# Improvements on Sample Preparation and Peptide Separation for Reduced Peptide Mapping Based Multi-Attribute Method analysis of Therapeutic Monoclonal Antibodies Using Lys-C Digestion

**DOI:** 10.1101/2022.02.28.482275

**Authors:** Xuanwen Li, Baibhav Rawal, Shannon Rivera, Simon Letarte, Douglas D. Richardson

## Abstract

The mass spectrometry based multi-attribute method (MAM) has gained popularity in the field of biopharmaceutical analysis as it promises a single method for comprehensive monitoring of multiple product quality attributes (PQAs) and product purity. Sample preparation for protein digestion and peptide separation are critical considerations for a reduced peptide mapping-based MAM. To avoid desalting steps required in tryptic protein digestion and in order to improve peptide separation for hydrophilic peptides, we developed an improved robust sample preparation using Lys-C protease for high-throughput MAM testing. Additionally, this method optimizes the peptide retention and separation of a stability-indicating VSNK peptide using a HSS T3 column for comprehensive PQA monitoring. A fully automated sample preparation had similar assay variations for PQAs monitoring compared to manual sample preparation. To the best of our knowledge, this is the first report of a high-resolution MS-based MAM using Lys-C digestion with enhanced PQA monitoring for hydrophilic peptides. The improved, robust MAM workflow for protein digestion and peptide separation will pave the way for broader MAM qualification and its applications for the characterization and quality control of therapeutic monoclonal antibodies.

## Introduction

Biotherapeutics, such as therapeutic monoclonal antibodies (mAbs) products, have become a predominant therapeutic modality for the treatment of a broad range of diseases in recent years. The global sales revenue for all mAb drugs totaled nearly $115.2 billion in 2018 and is expected to be valued at $300 billion by 2025.^1^ Most biotherapeutics are large, complex molecules or mixtures of molecules produced by living cells through multi-step bioprocess technology. Many naturally occurring post-translational modifications as well as chemically induced modifications are present in biotherapeutics during processing steps, manufacturing, and storage.^2-5^ Those product quality attributes (PQAs) include a variety of protein modifications, such as oxidation, deamidation, terminal variants and glycosylation. ^2-5^ Significant efforts have been made to understand the alterations incurred by these PQAs as well as the biological potency of therapeutic proteins after such modifications. Some product attributes may affect drug clearance, thereby altering drug efficacy. ^2-5^ Because of the complicated structure function relationship and heterogenous nature of some biotherapeutics, robust analytical platforms for ensuring consistent product quality during manufacturing and product storage through quality by design (QbD) approaches have started to gain popularity. ^2-3^

Multiple well-established analytical assays, including high performance liquid chromatography (HPLC) and capillary electrophoresis (CE) based methods are currently used for PQA monitoring and product purity testing throughout biotherapeutics development and production.^2-3^ If release/stability data from one of these assays falls outside of specification limits, orthogonal attribute assays like mass spectrometry (MS) assays can be employed to assist the root cause investigation. As regulatory requirements continue to evolve and expand, pharmaceutical industries are increasingly pressured to provide more detailed information on biologic products, while accelerating development timelines. ^2-3, 6^ Recently, MS-based methods intended to simultaneously monitor PQAs were introduced to assess the quality of biopharmaceuticals manufacturing processes in more detail. ^2-3, 6-8^ As such, the multi-attribute method (MAM), including PQA monitoring and purity testing through new peak detection (NPD), serve as an orthogonal method and could eventually replace several traditional HPLC and CE-based assays currently used for characterization and release of biopharmaceuticals.^9-19^

One of the challenges for the implementation of MAM in Good Manufacturing Practice (GMP)/ quality control (QC) areas is the streamlined sample preparation, which enables robust high-throughput testing while controlling assay variability. Reduced peptide mapping-based MAM (RPM-MAM) involves enzymatic digestion of proteins into peptides for MS detection.^20^ Trypsin or a trypsin/Lys-C mixture are the most frequently used proteolytic enzymes for high-resolution MS based MAM. ^9-19^ Besides its superior availability and affordability, trypsin digestion also offers some major advantages, such as an optimal average peptide length of ∼14 amino acids, ^21^ rendering tryptic peptides well-suited for MS analysis. Trypsin digestion efficiency, however, is less resistant to denaturants, such as guanidine hydrochloride, which is often used to denature therapeutic proteins.^21^ To improve the digestion efficiency and throughput, a desalting step carried out by buffer exchange is often used in the sample preparation of MAM.^9-15^ The extra desalting step brings challenges for high-throughput MAM testing with additional experimental variables, such as potential sample loss.^18^ The desalting steps increase the difficulty of fully automating sample preparation for MAM, although recent progress has been made with buffer-exchange plates and tips.^9-10, 13^ Additionally, trypsin is reported to have side activities, such as trace level of chymotrypsin activity, which may result in many unidentified peaks in the MAM.^22^ Collectively, the trypsin digestion efficiency, specificity and extra buffer exchange step bring additional complexity and variability for MAM.

Alternatively, lysyl endopeptidase (Lys-C) is considered specific and efficient in denaturing conditions. ^21^ Lys-C is relatively resistant to denaturants (such as 8 M urea), which enhances digestion efficiency and allows proteins to be digested in their optimal denatured states.^21^ The applications of using Lys-C digestion for high-resolution MS based MAM for PQA monitoring and NPD has not been reported. ^9-19^

Beyond sample preparation and protein digestion, peptide separation is another critical aspect of MAM development with the goal of capturing as many PQAs as possible. In the past, the changes observed on heat-stressed mAbs by ion exchange chromatography (IEX) were difficult to trace back to amino acid modifications by RPM. Recently, the root cause for changes in IEX profiles was linked to be the deamidation of the conserved heavy chain (HC) fragment crystallizable (Fc) VSNK peptide.^23-24^ A recent report indicates that the deamidation on VSNK has a functional impact on antibody-dependent cell-mediated cytotoxicity (ADCC) and FcγRIIIa binding for IgG1 mAbs.^24^ Deamidation of VSNK is often missed because this hydrophilic peptide is eluted near the void volume of most reverse phase (RP) columns after Lys-C or trypsin digestion.^25^ To capture this deamidation site, an additional chymotrypsin or Glu-C digestion has to be performed before LC-MS analysis.^23-24^ Optimization of LC conditions to detect the stability-indicating VSNK peptide and other hydrophilic peptides are critical for expanding the scope of RPM based MAM.

In this study, we first evaluated Lys-C and trypsin digestion efficiency for a therapeutic mAb in guanidine hydrochloride and optimized the separation of the VSNK hydrophilic peptide. The robustness of the improved MAM was evaluated to ensure its readiness for method qualification before transferring to QC labs.

## Materials and Methods

### Materials

Dithiothreitol (DTT) and iodoacetamide (IAM), both in no-weigh formats, were purchased from Thermo Pierce (Rockford, IL). Tris-HCl buffer (1 M, pH 8.0), guanidine HCl (8 M), water (Optima™), acetonitrile (ACN, Optima™), formic acid (FA), difluoroacetic acid (DFA), trifluoracetic acid (TFA), Zeba™ Spin Desalting Columns (7K MWCO, 0.5 mL) were purchased from Thermo Fisher Scientific (Waltham, MA). Ethylenediaminetetra-acetic acid (EDTA, 0.5 M, pH 8.0) and trypsin were purchased from Promega (Madison, WI). Lysyl endopeptidase (Lys-C) was purchased from Wako (Osaka, Japan). Endoproteinase Glu-C Sequencing Grade was purchased from Roche (Basel, Switzerland). Conductive-filtered CO-RE II tips (50-μL, 300-μL and 1-mL) and 50-mL reagent troughs, were purchased from Hamilton Company (Reno, NV). 96 deep-well plates (2-mL) were obtained from Eppendorf (Hamburg, Germany). Black Polystyrene Universal Microplate Lid was sourced from Corning (Corning, NY). 96-well Plate with Deactivated 700 µL Glass Inserts was purchased from Waters (Milford, MA). mAb-1 to mAb-8, were from the pipeline of Merck &Co. Inc. (Kenilworth, NJ, USA).

### Sample preparation and protein digestion

In-process or drug substance (DS) samples were diluted with water to 5 mg/mL. A total of 20 µL of the diluted sample (100 µg) was denatured and reduced in final solution (100 µL) containing 6 M Guanidine HCl, 50 µM Tris HCl, 50 µM EDTA and 200 µM DTT. The mixed sample was incubated in thermomixer at 37°C with 300 rpm shaking for 30 min. After mix and spin down, each sample was alkylated with 5 µL of 1M IAM protected from light at 25 00BAC for 30 min. A total of 5 μL of 200 mM DTT was added to block the unreacted IAM. Lys-C enzyme (1:10 (enzyme to mAb, wt: wt)) in 500 µL of 50 µM Tris-HCl (pH 8.0) was added to the protein samples with slowly pipette up/down 3 times to mix well. The digestion was incubated in a thermomixer at 37°C for 60 min. The digestion was quenched with 15 µL of 20% TFA. Digested samples were analyzed by LC-MS within 24 h after sample digestion. Otherwise, digested samples were stored at −80 ºC for future analysis. For the method optimization experiments with Lys-C or trypsin (1:10 (enzyme to DS, wt: wt)), the same preparation was used except aliquots (150 µL) were taken from the same digestion at the time point of 30 min, 60 min, 120 min and 240 min after adding the enzyme.

### UPLC-MS

Various reverse phase C18 columns, ion-pair reagents and concentrations for mobile phases, and column temperatures were screened for their retention of hydrophilic peptides. The columns evaluated included Acquity HSS T3 and BEH columns from Waters (Milford, MA), Zorbax 300 SB-C column from Agilent (Palo Alto, CA) and HALO ES column from Advanced Materials Technology (Wilmington, DE). All four columns in this evaluation were 2.1 × 150 mm in size.

For the final MAM, the LC was run with a Waters Acquity UPLC H-Class Bio system with 20 µL of sample injection per run onto the Waters HSS T3 column (100Å, 1.8 µm, 2.1 mm X 150 mm) with a column temperature of 40 ºC. Autosampler was set at 5°C. Mobile phase (MP)-A was 0.02% TFA in water and MP-B was 0.02% TFA in ACN. The flow rate was 0.3 ml/min. The LC gradient started with 0.1% MP-B from 0 to 5 min, 0.1%-12% MP-B from 5 to 7 min, followed by an increase to 40% MP-B over the next 38 min. The column was washed with 98% MP-B for 4 min before returning to 0.1% MP-B for balancing. The entire gradient run time was 1 hour. The divert valve from LC to MS was set at 2 min.

Eluted peptides were detected with a Q Exactive™ or Exactive Plus™ Orbitrap MS (Thermo) using Chromeleon (7.2.10, Thermo). The MS parameters for PQA monitoring included spray voltage at 3.8 kV, capillary temperature at 250 °C, sheath gas at 35 (arbitrary units), aux gas at 10 (arbitrary units), S-Lens RF level at 50, aux gas heater temperature at 250 °C, scan range from 300 to1800 m/z, AGC target at 1E6, maximum inject time at 200 ms, MS resolution at 140,000, mass acquisition time from 2.2 to 45.0 min, and lock mass 391.2843 m/z. For peak identification, the MS and MS/MS spectra were acquired in data-dependent acquisition (DDA) mode with the top 10 ions for fragmentation. The acquisition parameters were as follows: full MS resolution at 70,000, AGC target of 3E6, full MS scan range from 300 to 2000 m/z, full MS maximum injection time at 200 ms; dd-MS2 resolution at 17500, AGC target of 5E5, dd-MS2 maximum injection time at 100 ms, isolation window at 1.5 m/z, collision energy at 28, dynamic exclusion at 15 s.

### Robustness of MAM workflow

To evaluate the robustness of the MAM workflow, several assay variables in sample preparation were evaluated. For autosampler stability, the digested peptide preparations (n=6) were stored in the autosampler (5 °C) for up to 48 hours. The samples were run with LC-MS at times 0, 24 hr and 48 hr. For the Lys-C digestion time, the sample preparation was followed with the exception that a digestion time of 55 min (n=3) and 65 min (n=3) was utilized. For column temperature, the samples were run at 3 °C above (n=3) or 3 °C below (n=3) the target column temperature of 40 °C. To mimic sample transfer and aliquoting among different manufacturing and analytical testing locations, mAb-1 DS (n=3) was stored at −80 °C and thawed at room temperature for 20 min or until completely thawed, this freeze-thaw cycle was repeated 3 times. After 3 freeze-thaw cycles, samples were run with the MAM workflow at the same time as samples (n=3) without freeze-thaw treatment. To evaluate the stability of digested peptides at −80 °C in the event of any LC-MS failure and subsequent instrument downtime, digested peptides with different storage-day at −80 °C up to 28 days (4 weeks) were run by LC-MS at the same time. To evaluate mobile-phase stability, mobile-phases prepared day of analysis were compared to mobile-phases stored at room temperature for 4 weeks.

### Automated sample preparation and protein digestion with Lys-C

Automated Lys-C digestion was performed using the Hamilton Microlab STAR Liquid Handling System. The platform is configured with CO-RE 8-probe head designed to fit 50-μL, 300-μL and 1-mL tips. The Microlab STAR also accommodates iSWAP Robotic Transport Arm to move plates around the deck for incubation and reagents dispensing. The deck consists of four heater shaker units (HSUs) and one cold plate air cooled (CPAC) cooling unit.

The fully walk-away automated digestion method was developed in-house, and samples were digested in a 96-well plate which was protected from light with a dark lid except during reagent dispensing steps. The same mAb sample mixtures in the denaturing and reducing buffer as those in the manual digestion were transferred to an HSU maintained at 37°C and incubated for 30 mins while shaking at 300 rpm. A total of 5 µL of 1M IAM was added to the denatured/reduced samples and transferred to a 25ºC HSU for 30 mins. The alkylation reaction was quenched by adding 5 µL of buffer containing 1M DTT and 50 mM Tris-HCl buffer, pH 8.0. Lys-C enzyme (1:10 (enzyme to DS, w/w)) in 450 µL of 50 µM Tris-HCl (pH 8.0) were then added to the samples and incubated at 37°C for 60 mins. The digestion was quenched with 15 µL of 20% TFA and samples were analyzed immediately by LC-MS or stored at −80 ºC for future analysis.

### Lys-C/Trypsin and Glu-C digestion

mAb-8 samples were diluted to 5 mg/mL using LC-MS grade water. A total of 20 µL of diluted samples (100 µg) was reduced in 80 µL buffer containing 6.0 M guanidine hydrochloride, 50 mM Tris buffer (pH 8.0), 5 mM EDTA and 20 mM dithiothreitol (DTT) at 56°C for 30 min with shaking at 300 rpm. A total of 5 µL of 1 M IAM were added to the mixture and incubated at 25°C in dark for 30 mins; then 1 µL of 1 M DTT was added to quench the alkylation reaction. A total of 106 µL of the reaction mixture (100 µg mAb) was buffer exchanged into a solution containing 1M Urea, 50 mM Tris-HCl and 5 mM EDTA using Zeba Spin Desalting Columns 7K MWCO. The reaction mixture was then divided into two tubes (50 µL each). One 50 µL aliquot was used for Lys-C/Trypsin digestion, while the other for Glu-C digestion.

Lys-C/Trypsin Digestion: 50 µL of mAb-8 (50 µg), 100 μL digestion buffer (50 mM Tris buffer (pH 8.0)) and 2.5 μL of 0.5 mg/mL (1.25 μg) of Lys-C solution were added with an enzyme to substrate ratio of 1:40 (w/w) and incubated at 37 ºC for 1 hour with shaking at 300 rpm. Following Lys-C digestion, 2.5 μL of 0.5 mg/mL (1.25 μg) of trypsin were added to the sample mixture with an enzyme to substrate ratio of 1:40 (w/w) and incubated at 37 ºC for 3 hours. Finally, 10 μL of 20% TFA were added to the samples to quench the reaction.

Glu-C Digestion: 50 µL of mAb-8 (50 µg), 100 μL digestion buffer (50 mM Tris buffer (pH 8.0)) and 5 μL of 0.5 mg/mL (2.5 μg) of Glu-C solution were added using an enzyme to substrate ratio of 1:20 (w/w) and incubated at 37 ºC for 4 hours with shaking at 300 rpm. Following digestion, 10 μL of 20% TFA were added to the samples to quench the reaction. The digested samples were analyzed immediately by LC-MS or stored at −80 ºC for future analysis.

### Data analysis

For mAb characterization during method development and optimization, the MS data with MS/MS acquisition were searched against BioPharma Finder 3.2 (Thermo, Waltham, MA). To evaluate the digestion efficiency, ‘RelativeLoad’ and ‘PeptideMapQuality’ in BioPharma Finder were used. ‘RelativeLoad’ displays a measure of the protein quantification using the top 3 peptides normalized to 100% of the best digested sample among all the samples submitted for analysis. ‘PeptideMapQuality’ displays a measure of the quality of the digestion. A value of 1 indicates the perfect digestion that the peptides in the sample are neither under-digested nor over-digested.

For PQA quantification in Chromeleon (Thermo, Waltham, MA), the two or three most abundant charge states (each with the top two or three most abundant isotopes) from the method development studies were selected for data processing and reporting. The PQA list included two asparagine (N) deamidations, one methionine (M) oxidation, one heavy chain N-terminal variant (HC-pE), and two heavy chain C-terminal variants (HC+K and HC-L-amidation). The mass tolerance was set at 5 ppm. The extracted ion chromatograms (EICs) of non-modified and modified peptides were used to determine the percent of PTMs (Post Translational Modifications) using the formula [peak area of the extracted ion chromatogram (EIC) of the modified peptide] / [peak area of EIC of the modified peptide + peak area of EIC of the unmodified peptide] × 100.

## Results

### Sample preparation optimization for protein digestion

To drive potential high-throughput MAM testing and avoid desalting steps during sample preparation, a time-course experiment was performed to evaluate the digestion efficiency with both Lys-C and trypsin. Surprisingly, Lys-C was able to achieve almost comparable digestion within 30 min as indicated by the number of MS peaks for heavy chain and light chain compared to those in 240 min digests (Figure 1A). Both the heavy chain and light chain had sequence coverage above 99% (Table 1 and supplementary Figure 1). The number of MS peaks labelled as unidentified from BioPharma Finder remains relatively constant up to 240 min (Figure 1A). For the commonly used trypsin used for RPM, the number of identified MS peaks for the heavy chain and light chain increased with digestion time, and the number of unidentified peaks decreased (Figure 1B). The lower efficiency of trypsin digestion in denaturing buffer was indicated by ‘RelativeLoad’ and ‘PeptideMapQuality’ calculations in Biopharma Finder (Figure 1C and 1D). Both ‘RelativeLoad’ and ‘PeptideMapQuality’ for trypsin digestion increased over digestion time, and the ‘PeptideMapQuality’ was still below 0.2 up to 240 min of digestion time (Figure 1C). In contrast, samples from Lys-C digestion remain close to 100% for ‘RelativeLoad.’ ‘PeptideMapQuality’ was close to 1 starting at 30 min of digestion time, indicating that the peptides in the sample were neither under-digested nor over-digested with Lys-C digestion (Figure 1D). Consistent with the observations from BioPharma Finder analysis, the TIC (total ion chromatogram) profiles of digested peptides with Lys-C were similar from 30 min to 240 min (Figure 2).

**Table 1.** Sequence coverage of seven mAbs with the improved MAM workflow.

| mAb | Sequence Coverage |  |
| --- | --- | --- |
|  | HC | LC |
| mAb-1 | 99.1% | 99.1% |
| mAb-2 | 98.9% | 99.1% |
| mAb-3 | 98.9% | 99.1% |
| mAb-4 | 98.6% | 99.1% |
| mAb-5 | 98.9% | 97.7% |
| mAb-6 | 98.9% | 99.1% |
| mAb-7 | 98.2% | 99.1% |

Seven mAbs were prepared and run with the optimized MAM workflow with MS/MS an acquisition. Sequence coverages were obtained from Biopharma Finder. HC: Heavy chain; LC: light chain

**Figure 1:**
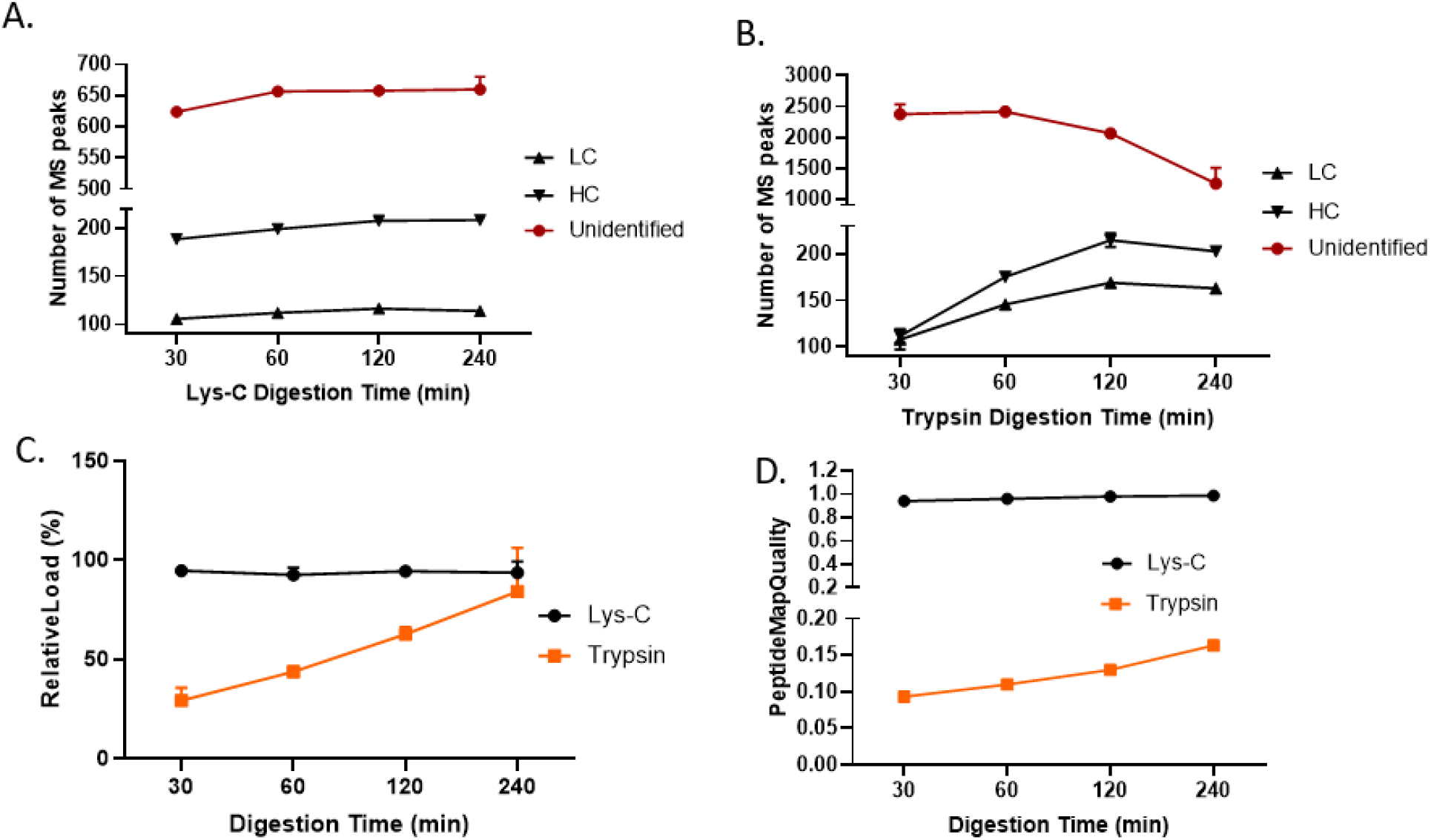
The time-course study of Lys-C and Trypsin digestion. mAb-1 samples (n=3) were digested with Lys-C or trypsin (1:10) according to the sample preparation procedures. Aliquots (150 µL) from the same digestion sample was taken at 30 min, 60 min, 120 min and 240 min for LC-MS/MS analysis. The data analysis was run with Biopharma Finder. The number of MS peaks for mAb-1 HC, LC, unidentified peaks, ‘RelativeLoad’ and ‘PeptideMapQuality’ in BioPharma Finder were used to compare digestion efficiency with both enzymes. LC: light chain; HC: heavy chain.

**Figure 2:**
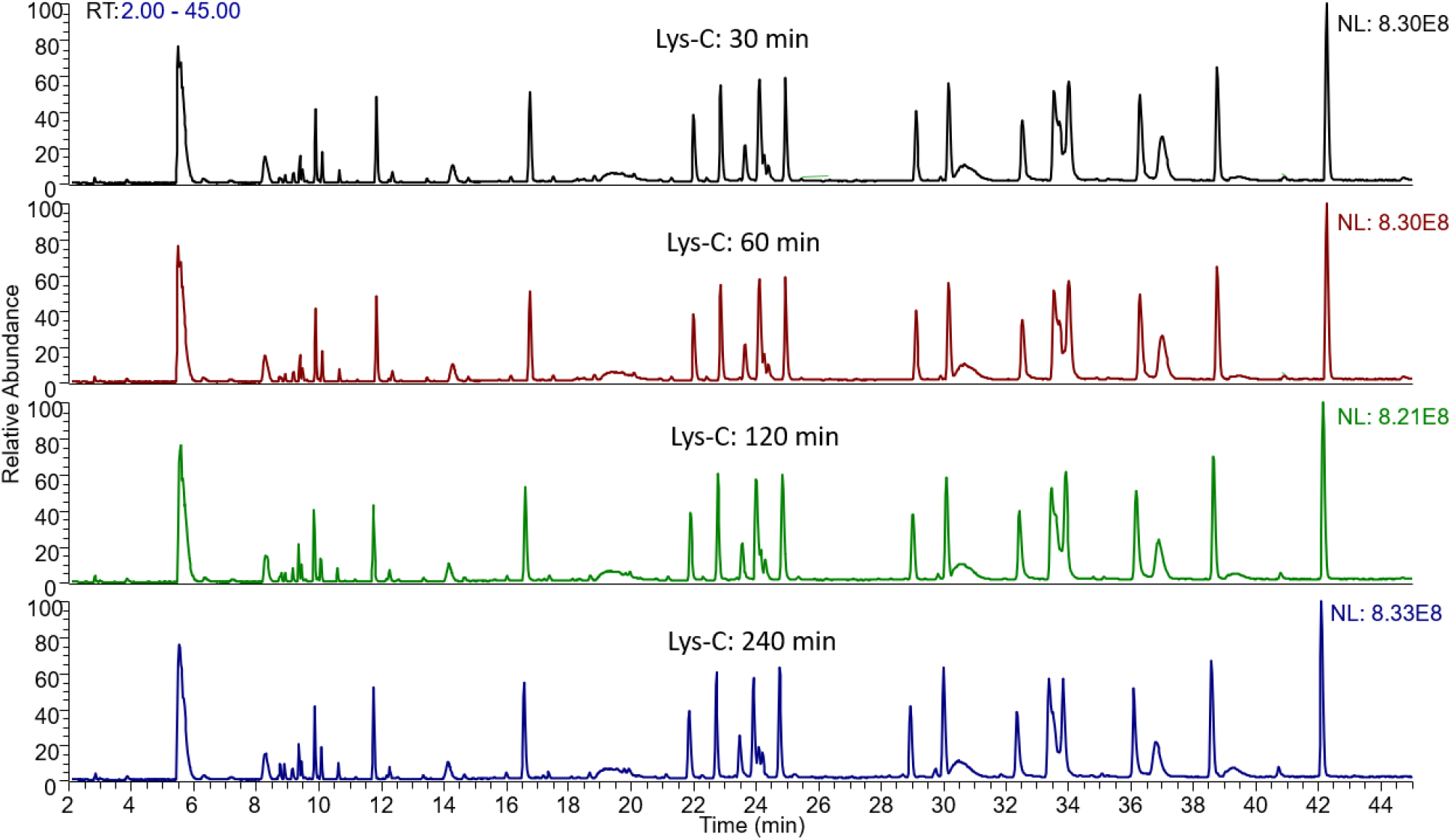
The TIC of digested peptides from the time-course study of Lys-C digestion. mAb-1 samples were digested with Lys-C (1:10) according to the sample preparation procedures. Aliquots (150 µL) from the same digestion sample was taken at 30 min, 60 min, 120 min and 240 min for LC-MS/MS analysis.

To select the most suitable time point for Lys-C digestion, the levels of certain PQAs were monitored. Similar to the number of MS peaks identified from heavy chain and light chain (Figure 1A), the levels of most PQAs, including heavy chain N and C terminal variants, and oxidation of a methionine residue at the constant Fc region, remained relatively stable from 30 to 240 min (Figure 3). Asparagine (Asn) deamidation and succinimidation, however, showed time-dependent increase with digestion time as represented by the well-known PENNYK peptide (Figure 3). To ensure robustness of sample preparation for digestion time, and testing logistics of the sample process and LC-MS readiness for the entire MAM workflow, the digestion time of 60 min for Lys-C was selected for the final method.

**Figure 3:**
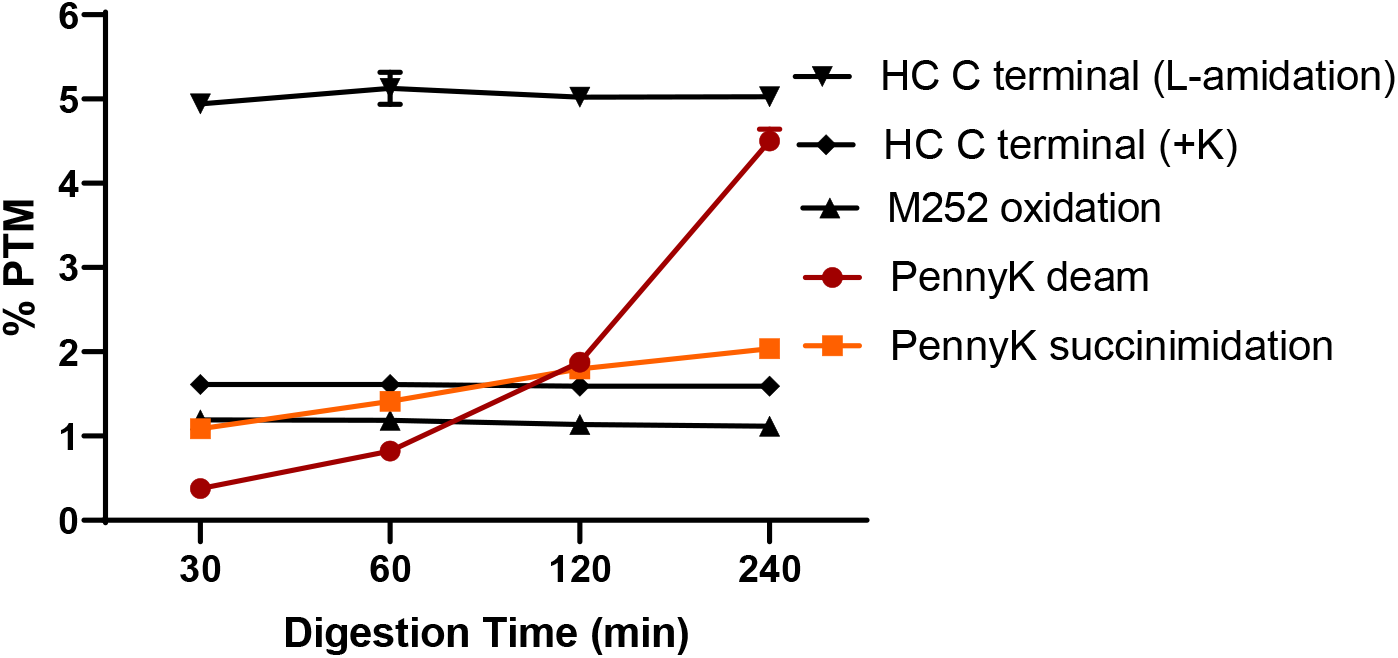
The %PTM of several PQAs from Lys-C digestion. mAb-1 samples were digested with Lys-C (1:10) according to the sample preparation procedures (n=3). Aliquots (150 µL) from the same digestion sample was taken at 30 min, 60 min, 120 min and 240 min for LC-MS/MS analysis. %PTM of PQAs were quantified with Chromeleon.

Lys-C protein digestion offers a universal sample preparation method for mAb digestion. The sequence coverages of seven different single mAbs and three mAb co-formulations (different ratios of two different mAbs) with the same sample preparation using Lys-C digestion were over 96.5% for both heavy chain and light chain (Table 1 and Table 2).

**Table 2.** Sequence coverages of 3 mAb coformulations with the improved MAM workflow.

| mAb<br>coformulation | Sequence Coverage |  |  |  |
| --- | --- | --- | --- | --- |
|  | mAb-1/HC | mAb-1/LC | Co-HC | Co-LC |
| mAb-1/mAb-2 | 99.1% | 99.1% | 98.9% | 99.1% |
| mAb-1/mAb-3 | 99.1% | 99.1% | 99.3% | 99.1% |
| mAb-1/mAb-4 | 99.1% | 99.1% | 96.8% | 97.7% |

Three mAb coformulations with different ratios of mAb-1 to 3 coformulated mAbs (mAb-2, mAb-3 and mAb-4) were prepared and run with the optimized MAM workflow with MS/MS acquisition. Sequence coverages were obtained from Biopharma Finder. HC: Heavy chain; LC: light chain; Co-HC: heavy chain from the co-formulated mAb; Co-LC: light chain from the co-formulated mAb.

### Optimization of UPLC separation for hydrophilic peptides

To develop a comprehensive MAM to capture all PQAs and new peaks, several UPLC conditions, such as different column chemistry, column temperatures, ion-pairing reagents, and their concentrations, were screened for their ability to capture hydrophilic peptides by using VSNK as the model peptide. The void volume for the columns used (2.1 × 150 mm) was determined to be approximately 1.6 min with the flow rate at 0.3 ml/min measured from the UV profile with a negative peak from injection of a blank (data not shown). A 2-min divert valve for LC from waste to MS detection was selected to ensure enough desalting for robust LC-MS setup. As shown in Figure 4, HSS T3 column was ideally suited for the enhanced retention of hydrophilic VSNK peptide compared to the BEH column. For mobile phase additive, the latest retention time of VSNK peptide retention is 0.1% TFA> 0.02% TFA>0.1%DFA. The VSNK peptide cannot be captured with 0.1% FA with the 2-min divert switch. Additionally, lower column temperature helped retain hydrophilic VSNK peptide. Both Zorbax and Halo columns failed to retain the VSNK peptide with 0.02% TFA and column temperature at 45 °C with the 2-min divert switch (data not shown).

**Figure 4:**
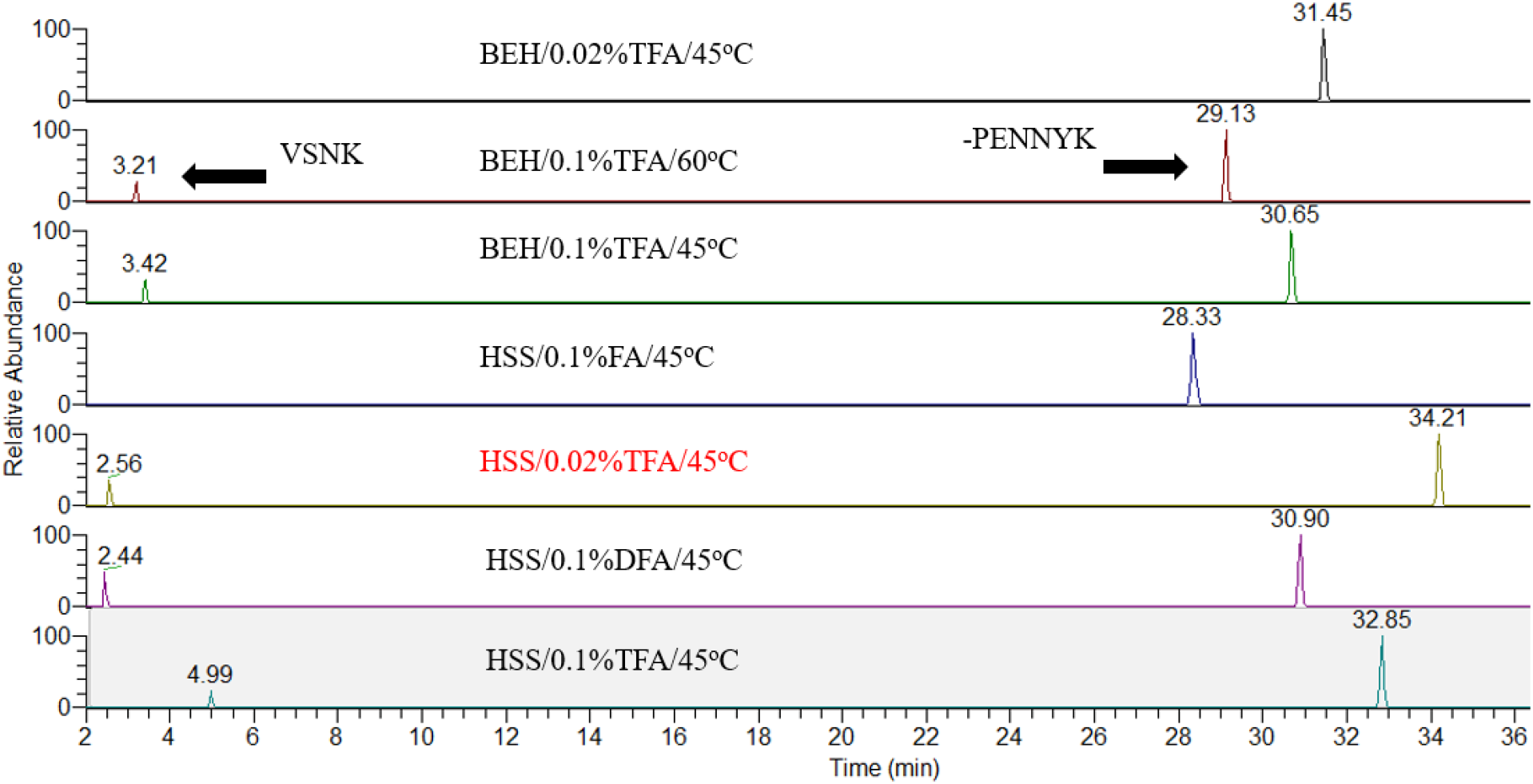
The optimization of peptide retention of the VSNK peptide. Digested mAb-1 samples were run with different column, ion-pair reagents and concentrations, and column temperatures. The void volume for the column used (2.1 × 150 mm) was determined to be at approximately 1.6 min with the flow rate at 0.3 ml/min as measured from the UV profile with a negative peak from injection of a blank. The 2-min divert valve from LC to MS was selected to ensure enough desalting for robust LC-MS setup. The extracted ion chromatogram (EIC) of the VSNK and PENNYK peptides were visualized for comparison.

To drive the limit of quantification of PQA monitoring, MS sensitivity is another consideration for the entire MAM workflow. The MS intensity of several peptides, including the VSNK peptide, showed about 3-fold higher with 0.02% TFA compared to 0.1% TFA with the HSS T3 column (Figure 5). Additionally, according to the manufacture guidelines, temperatures between 20–45 °C are recommended for operating HSS columns in order to enhance selectivity, lower solvent viscosity, and increase mass transfer rates. Taken together, the HSS T3 column with 0.02% TFA and column temperature 40 °C was selected for the MAM, which provided balanced retention of hydrophilic peptides and MS intensity, and avoided the upper limit of column temperature 45 °C.

**Figure 5:**
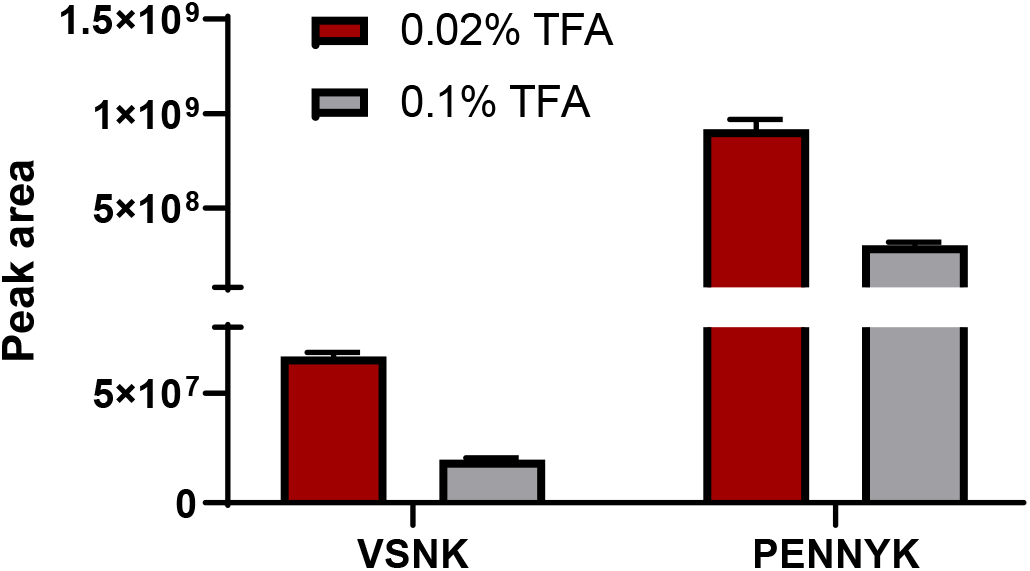
The MS responses of the VSNK and PENNYK peptides with different mobile phase additives. Digested mAb-1 samples were run with the HSS T3 column with 0.1% or 0.02% TFA (n=7). The peak areas of the non-modified VSNK and PENNYK peptides were compared.

With the selected final sample separation conditions, VSNK and its deamidated forms, which have slightly later elution times compared to unmodified VSNK, were well separated and resolved (Figure 6A). The peak IDs were confirmed with accurate mass (Figure 6B and 6C) and MS/MS (data not shown). The PENNYK peptide and its deamidated species were also well resolved with the HSS T3 column (data not shown). The deamidation levels of VSNK from Lys-C/trypsin digested samples (the same VSNK as that from Lys-C digestion) using the LC-MS conditions of the established MAM workflow were highly consistent with the same samples digested with Glu-C, a less frequently used enzyme (Figure 7). Glu-C digestion resulted in a longer peptide for better peptide retention and separation as reported before.^24^ The increase of deamidation on VSNK in mAb-8 from real-time stability samples using both digestion methods further demonstrated the importance of this modification, and it was the only modification correlating with the changes of IEX profiles in those samples (data not shown).

**Figure 6:**
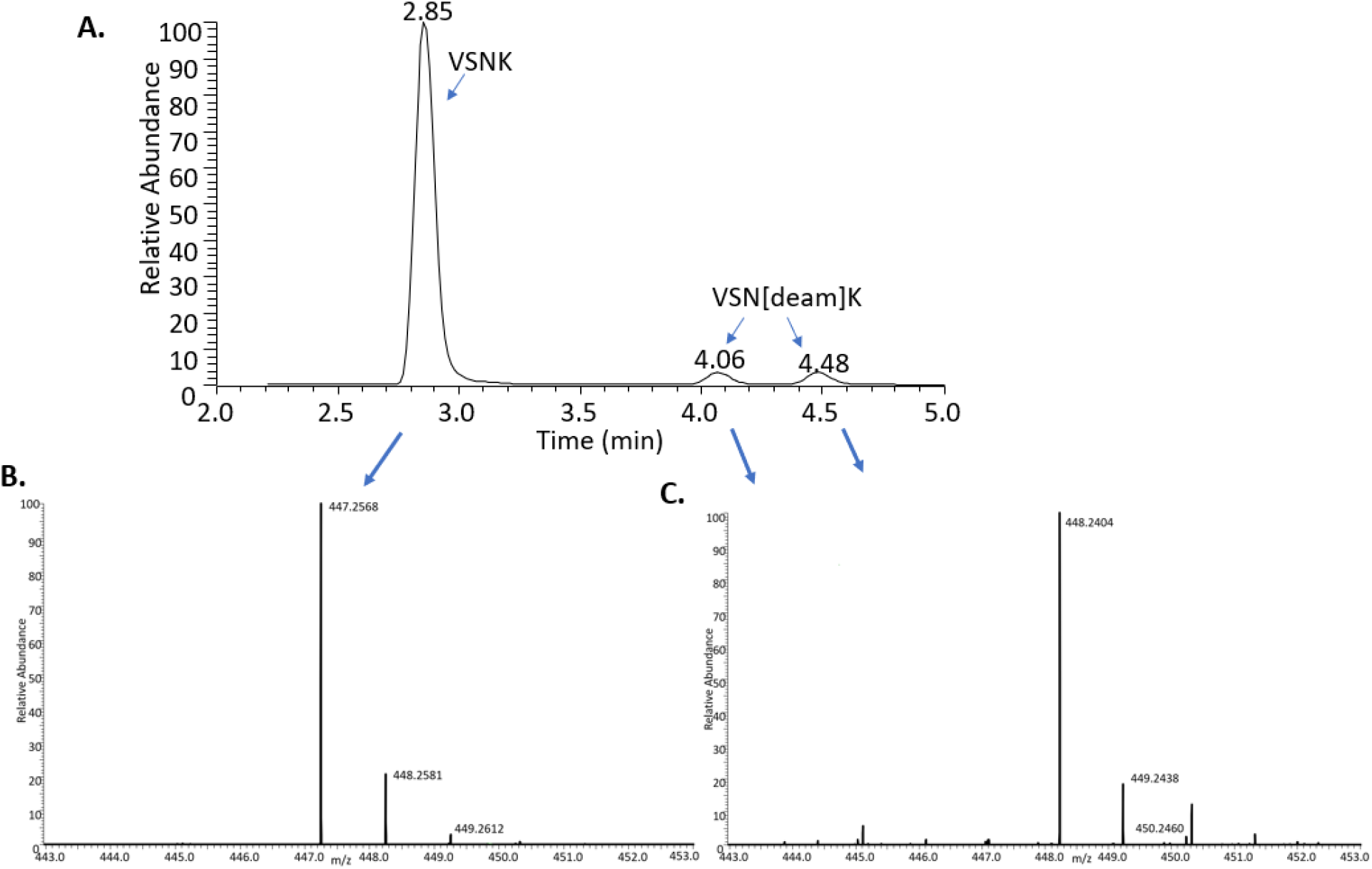
The separation of the VSNK peptide and its deamidated forms from heat-stress mAb-1. Heat-stress mAb-1 samples (50 °C/day 7) were run with the HSS T3 column with 0.02% TFA. The EIC (A) and accurate mass of the non-modified (B) and deamidated VSNK (C) were analyzed.

**Figure 7:**
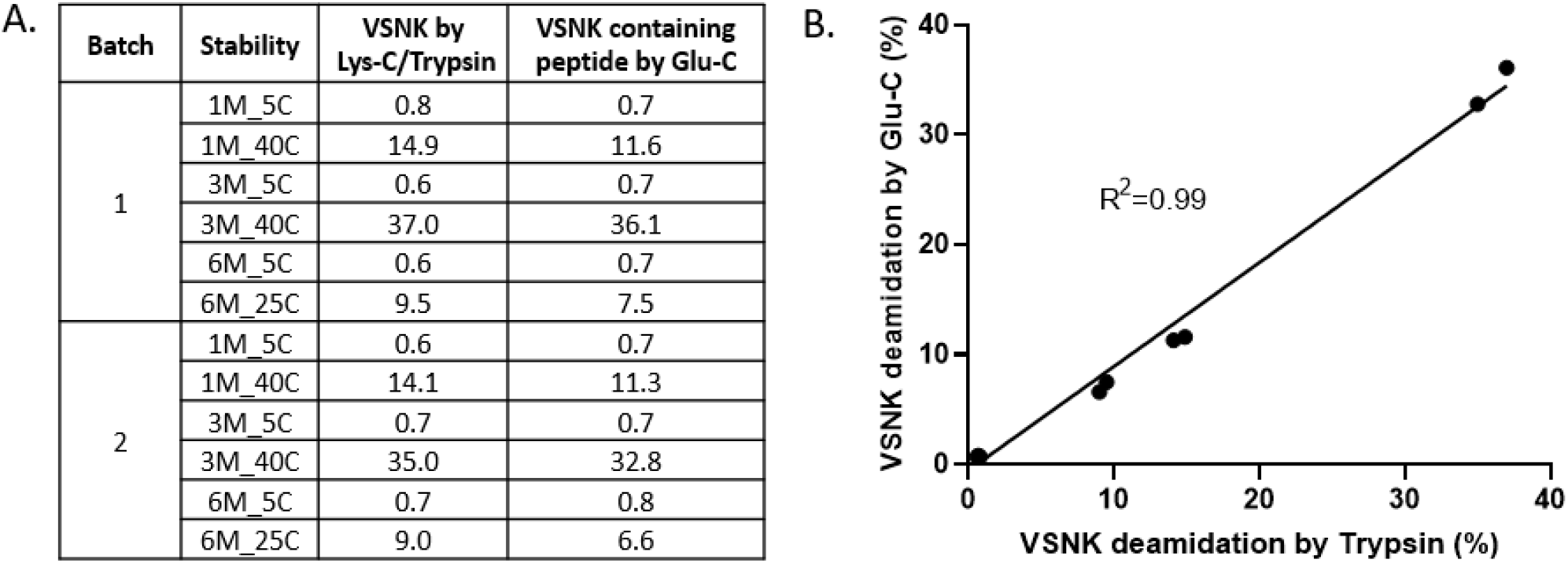
The comparison of deamidation of VSNK from trypsin and Glu-C. Real-time stability samples from 2 batches of mAb-8 were digested with Lys-C/trypsin or Glu-C. The VSNK peptide from trypsin digestion were run with the LC-MS conditions in the improved MAM. The deamidation levels were compared to a Glu-C digestion designed to specifically capture this critical VSNK deamidation.

### Robustness

To evaluate the robustness of the MAM workflow, several assay variables in sample preparation, including autosampler stability up to 48 h, Lys-C digestion time, column temperature, freeze-thaw stability of both protein and digested peptides, and mobile phase stability were evaluated. All PQAs remained at similar levels despite variations in all parameters tested except for deamidated peptides in relation to autosampler stability (supplementary Figure 2, 3 and 4). There was a clear increase in deamidation levels with time in the autosampler (supplementary Figure 2A and 2B). The absolute changes, however, were less than 1% for both deamidation sites up to 48-hour in the autosampler, and the final deamidation levels for both deamidation sites were low up to 48-hour in the autosampler. Both oxidation and deamidation levels increased after 48-hour in the autosampler (data not shown). The impact of autosampler incubation on PQAs was considered minimal if the samples were tested within 24 h between completion of sample preparation and LC-MS data acquisition of the last sample. Otherwise, the digested samples are recommended to be stored in a −80 °C freezer. The one cycle of freeze-thaw stability of digested peptides in a freezer for up to 4 weeks had no detected changes in the stability of all PQAs (Figure 4).

### Automated sample preparation

To drive consistency in sample preparation across different analytical labs and the throughput of MAM testing, the automated sample preparation using the Hamilton STAR system was tested. The automated preparation showed similar (over 99%) sequence coverage for both heavy chain and light chain as consistent with manual digestion (data not shown). The PQAs had similar PTM levels and assay variations in different days compared to those from the manual preparation (Figure 8).

**Figure 8:**
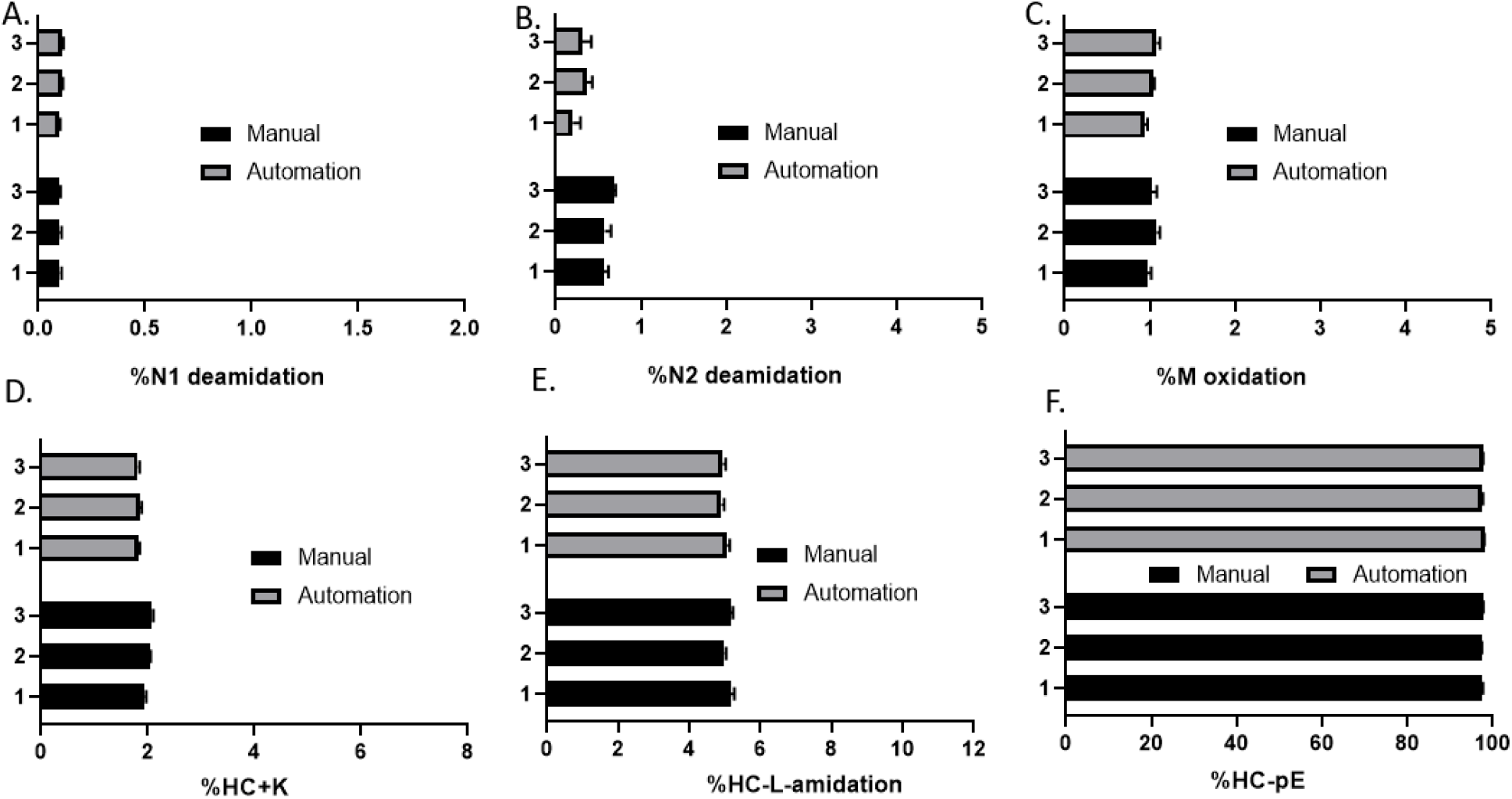
The comparison of PTM levels of PQAs between automated and manual protein digestion. mAb-1 DS samples were digested with manual or automated digestion methods over three different days (n=6 for each day). The PTM levels of PQAs were compared. The y-axis means three different days.

## Discussion

A MAM with improved sample preparation using Lys-C digestion and peptide separation was developed. The optimized MAM was demonstrated to be robust and comprehensive for PQA monitoring.

### Sample preparation and digestion strategies of MAM

The efficiency, specificity and throughput of sample preparation are among the key considerations for a streamlined MAM workflow. Trypsin is the most widely used enzyme for RPM-MAM using high-resolution MS in the literature thus far. ^9-15^ The time-course evaluation of trypsin digestion highlights the need of a desalting step for sample preparation in order to achieve fast and complete digestion (Figure 1).^10, 15, 20^ The desalting step in a trypsin digestion may bring another source of variations and prevent high-throughput MAM testing and the full automation of MAM due to the many challenges of implementing desalting steps in an automated workflow. In additional, the desalting tips, plates or spin columns needed for both manual and automatic sample preparation may encounter supply chain issues for timely MAM release testing in commercial quality control labs. Lys-C overcomes these challenges compared to trypsin. The advantages include 1) a nearly complete digest in denaturing buffer as short as 30 min (Figure 1); and 2) less non-specific digested and missed cleavages peptides (Figure 1). The lower number of peptides after digestion, especially those unidentified peptides, were considered beneficial for NPD of MAM (manuscript in preparation).

One of the challenges for Lys-C digestion is that the enzyme generates longer peptides compared to trypsin. For example, the peptide that reports on methionine oxidation in our MAM workbook has 40 amino acids. With a high-resolution mass detector, there is minimal technical challenge to specifically monitor this peptide and its modified form. However, we did observe challenges for generating high-quality MS/MS data using higher collision-energy dissociation (HCD) in a Q Exactive system for such long peptides to confirm the exact modification sites. Trypsin may still be the best option during method development to fully understand the degradation pathway of the peptide before switching to Lys-C digestion for PQA monitoring and NPD. Xu et al reported a peptide mapping method using Lys-C with a Quadrupole Dalton (QDa) mass detector to selectively monitor and quantitate PTMs in a therapeutic monoclonal antibody. The data from the low-resolution QDa based method provided comparable results to multiple traditional assays.^26^ We anticipate that high-resolution MS based MAM will bring additional specificity for certain PQAs and drive the limit of quantification with the same digestion method using Lys-C.

### Peptide separation of MAM

The ability of RPM-MAM to monitor multiple attributes in a single assay offers advantages over traditional HPLC or CE based assays with higher level of sensitivity and specificity using a high-resolution MS. Some critical attributes may be missed when reverse phase HPLC is used for peptide separation before MS analysis, because certain amino acids are located on short, hydrophilic peptides that are not typically retained. Among all protein modifications, asparagine deamidation is a major degradation pathway for therapeutic mAbs.^4-5^ Deamidation in the constant Fc region which is detected on the short peptide VSNK leads to higher acidic fractions in IEX based assay,^23-24^ which was reported to impact mAb Fc-effector function, including ADCC.^24^ This critical deamidation site is often detected only on missed cleavage peptide resulting in a larger peptide or often undetected completely by Lys-C or trypsin-based reduced peptide mapping because of the hydrophilic nature of the VSNK peptide. ^16-17, 25^ For example, VSNK was one of the few undetected peptides from an optimized LC-MS/MS peptide mapping protocol for NIST mAb with close to 97% sequence coverage.^25^ Additional protein digestion with chymotrypsin or Glu-C was needed to detect this site. ^23-24^ Lu et al developed a tryptic peptide mapping assay with a BEH C18 column, which could detect and accurately quantify the deamidation on VSNK comparable to a Glu-C digestion method.^24^ However, the mobile phase conditions were not disclosed. The VSNK peptide was not captured by a BEH column with a 2-min divert at 0.3 ml/min using our final mobile phase conditions (0.02%TFA and column temperature 40°C) as shown in Figure 4. In contrast, the HSS T3 C18 column was able to capture the VSNK peptide far outside of the void volume (Figure 4), which indicates that the HSS T3 C18 column enables a robust LC-MS system with longer time for desalting and less frequent MS source cleaning. The retention of the VSNK peptide was enabled by the complex interaction of the selected stationary phase, concentration and type of mobile phase additives and column temperature. The ion pairing reagent 0.1%TFA provided the best retention of the VSNK peptide (Figure 4). The chromatographic separation of peptides is influenced by not only the mobile phase composition, but also the MS performance (e.g., sensitivity, charge state and adduct ions), which is also strongly affected by the mobile phase composition. The final phase conditions (0.02%TFA and column temperature 40°C) were considered to be the right balance of peptide separation and MS performance.

The optimized LC-MS conditions designed to retain VSNK peptide removes the need for extra Glu-C digestion for both trypsin and the Lys-C-based MAM workflows. The improved LC conditions may serve as a platform method for reduced peptide mapping to capture all potential mAb PTMs in a single assay to understand the structure-function relationship of mAbs.

### Robustness and automation of MAM

With the improved sample preparation and peptide separation, the MAM for PQA monitoring was considered relatively robust. Several variables of the MAM, including autosampler stability up to 24 h, Lys-C digestion time, column temperature, freeze-thaw stability of protein and digested peptides, and mobile phase stability were evaluated to have minimal impact on PQA levels. Future efforts will include additional variables of the MAM using DOE to provide a better understanding of the assay performance for MAM validation.

The combined MAM workflow can be completed within 4 hours (1 hour for sample denaturation, reduction and alkylation, 1 hour for Lys-C digestion and 1 hour for LC-MS) with the testing throughput for nearly 20 samples in a single day (24 hours) using manual digestion (considering additional time for samples used for system suitability testing). As mentioned earlier, Lys-C protein digestion offers a universal sample preparation method for mAb digestion with the potential for full automation of sample preparation without desalting steps. Our initial evaluation of the full automation of sample preparation with Lys-C digestion using an automated liquid handler showed very comparable levels of several PQAs and assay precisions compared to the manual digestion (Figure 8). The throughput of sample preparation using automation system can be increased dramatically, however, the throughput of sample preparation will outpace LC-MS analysis because of autosampler stability.

The MAM performance for identity, PQA monitoring and NPD using the developed sample preparation and peptide separation method was fully qualified (manuscript in preparation). The qualification results ensured assay performance before transferring it to a QC environment.

## Conclusions

In this study, we developed an improved robust MAM workflow for sample preparation and peptide separation. Lys-C digestion without additional desalting steps increased the throughput of MAM testing and allowed for full automation of sample preparation. The HSS T3 column and UPLC mobile phase conditions increased the likelihood of capturing the hydrophilic VSNK peptide to expand the coverage of the MAM. The improved MAM workflow is the foundation for a platform-based approach for the use of high-resolution based MAM for the characterization and quality control of therapeutic mAbs.

## Acknowledgments

We thank the members of the Analytical R&D Mass Spectrometry, and the internal MAM working group for fruitful discussions. The feedback and early contributions for the method development from Yan-Hui Liu, Wang Yi, Haitao Jiang, Rong-Sheng Yang, Michael Olma, Bhumit Patel, Brent Kochert, Xinliu Gao, Bernard Choi and Zachary Dunn are highly appreciated. We thank Richard Rogers and Yuko Ogata from Just Biotherapeutics for early discussions during MAM development. We thank Hillary A. Schuessler and Brent Kochert for critical review and edits of the manuscript.

## Supplementary Figures

**Supplementary Figure 1.**
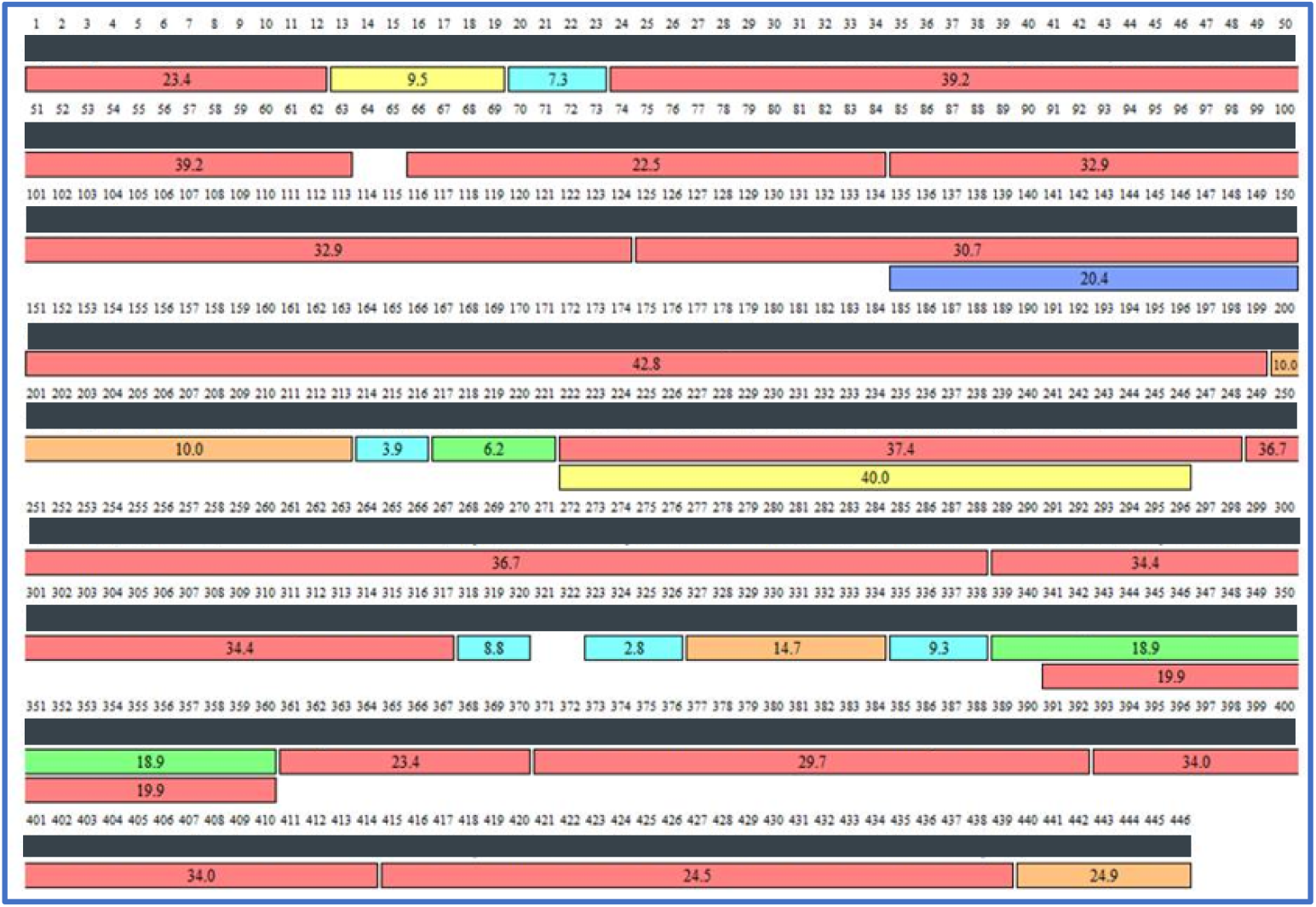
The sequence coverage of heavy chain from mAb-1 with the optimized Lys-C digestion. Over 99% of sequence coverage was detected from BioPharma Finder analysis. Minimum miss-cleavage peptides were produced from Lys-C digestion. The proprietary amino acid sequences were intentionally covered with black bar.

**Supplementary Figure 2.**
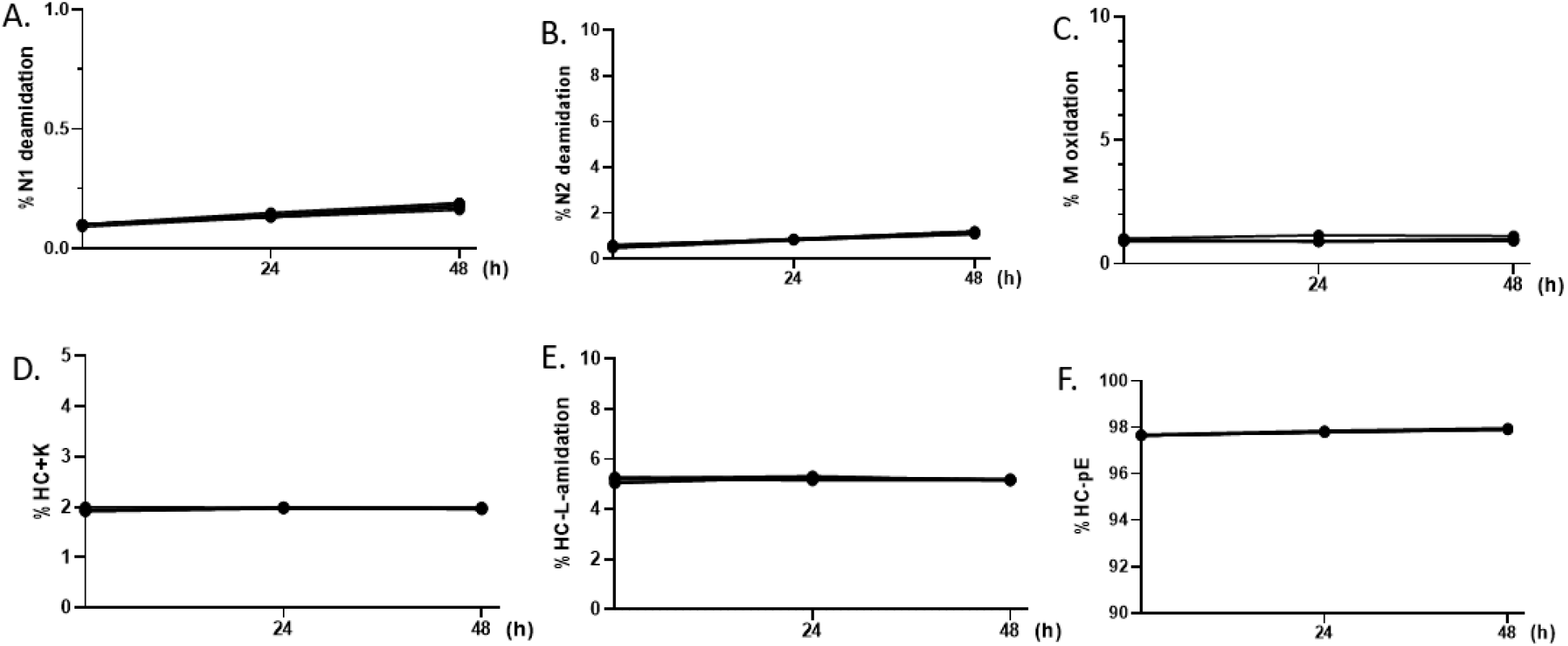
The autosampler stability of 6 PQAs from mAb-1. The digested peptides (n=6) from mAb-1 were stored in the autosampler (5°C) for up to 48 hours. The samples were run with LC-MS at time 0, 24h and 48h. The stability of %PTMs of PQAs from Chromeleon were compared.

**Supplementary Figure 3.**
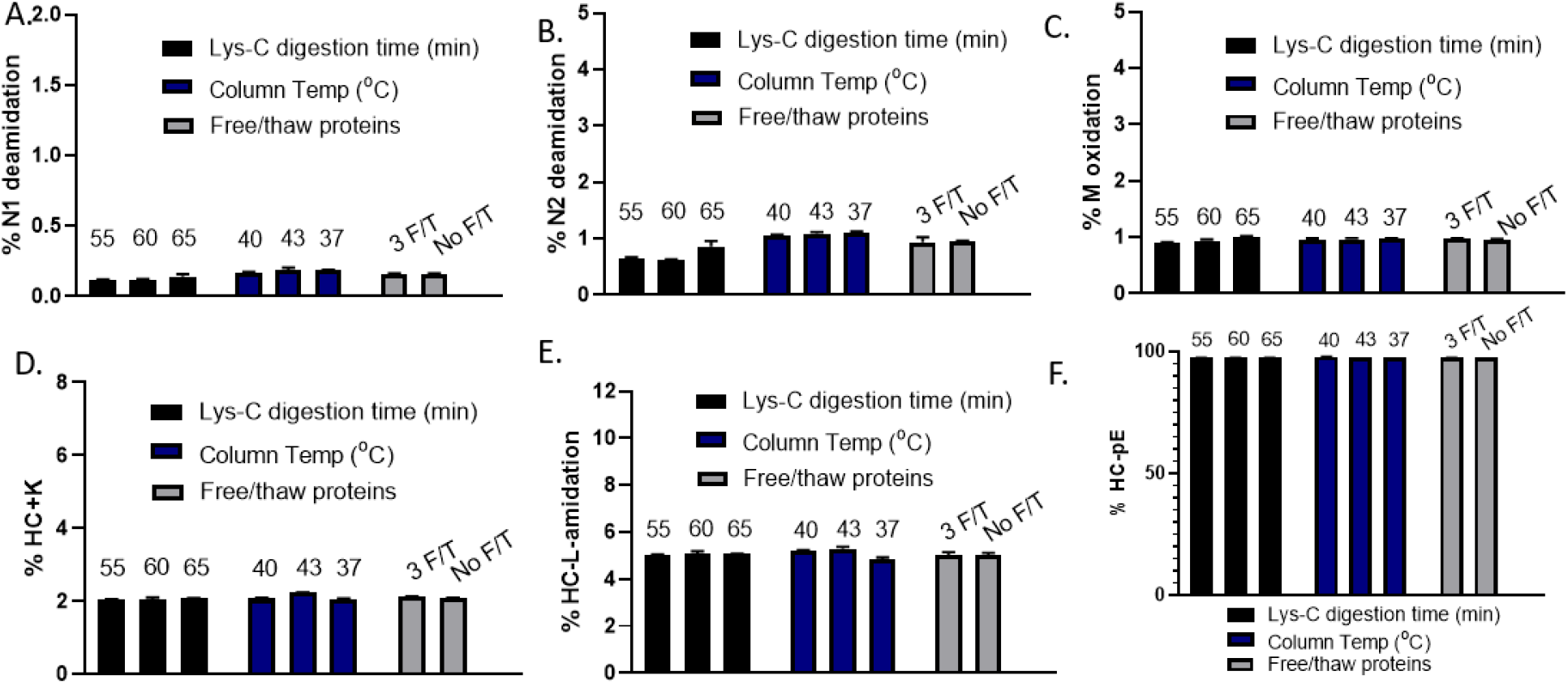
The robustness of Lys-C digestion time, column temperature and freeze/thaw of mAb proteins. For Lys-C digestion time, the sample preparation was followed except for some samples (n=3) were digested with 55 min or 65 min. For column temperature, the same samples (n=3) were run with 37 °C, 40 °C or 43 °C. For freeze/thaw stability, mAb-1 proteins (n=3) went through 3-cycles of freeze-thaw. The stability of %PTMs of PQAs from Chromeleon were compared.

**Supplementary Figure 4.**
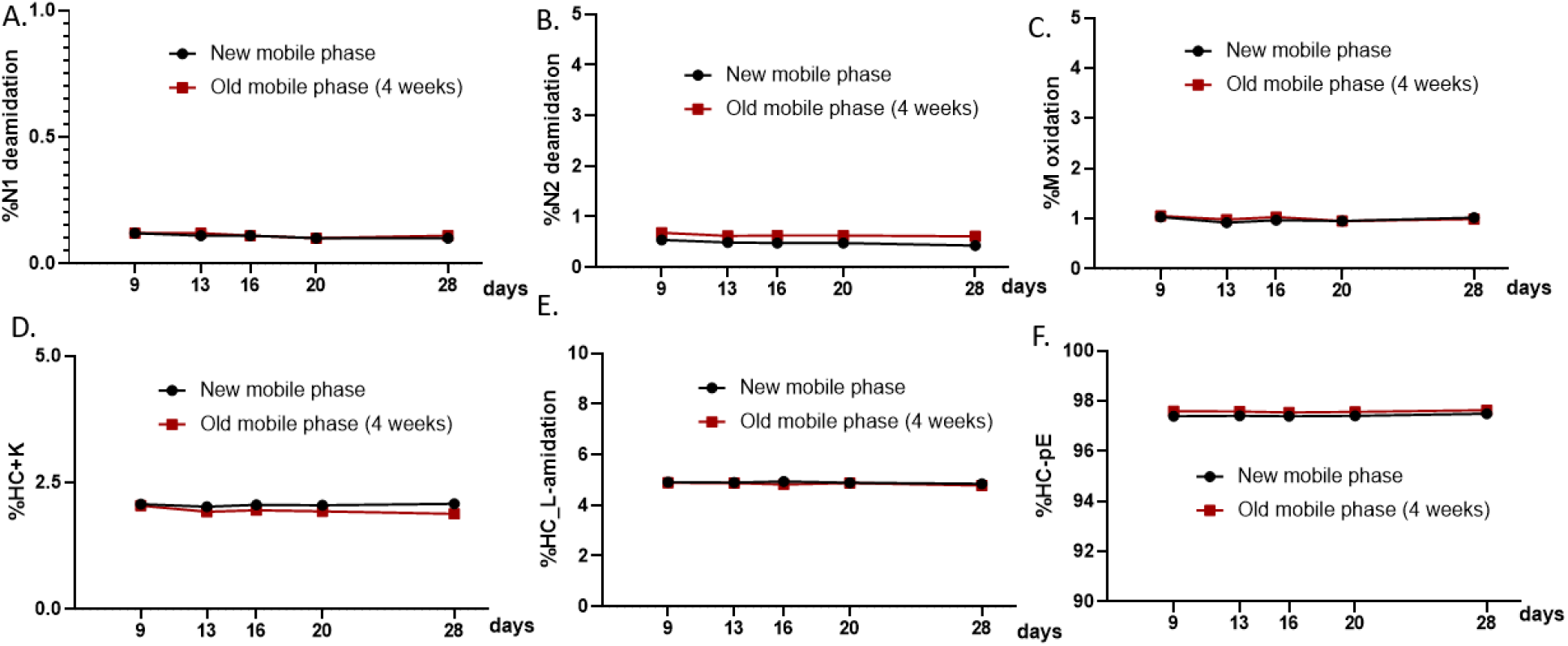
The stability of digested peptides and LC mobile phases. Digested peptides with different storage-day in −80 °C up to 28 days (4 weeks) were analyzed via LC-MS with fresh mobile phases or old mobile phase with 0.02%TFA stored at room temperature for 4 weeks. The stability of %PTMs of PQAs from Chromeleon were compared.

## Notes

### Competing Interest Statement

The authors have declared no competing interest.

